# Nectriatides with no antifungal activity bind to ergosterol and potentiate the antifungal activity of amphotericin B against *Candida albicans*

**DOI:** 10.1101/2025.02.04.636370

**Authors:** Keisuke Kobayashi, Ryosuke Miyake, Kenichiro Nagai, Yukino Sato, Reiko Seki, Kumiko Sakai-Kato, Shinichi Nishimura, Taichi Ohshiro, Hiroshi Tomoda

**Affiliations:** Department of Microbial Chemistry, Graduate School of Pharmaceutical Sciences, Kitasato University, Tokyo, Japan; Medicinal Research Laboratories, School of Pharmacy, Kitasato University, Tokyo, Japan; Laboratory of Analytical Chemistry, School of Pharmacy, Kitasato University, Tokyo, Japan; Department of Biotechnology, Graduate School of Agricultural and Life Sciences, The University of Tokyo, Tokyo, Japan; Graduate School of Integrated Sciences for Life, Hiroshima University, Higashi-Hiroshima, Japan

**Keywords:** nectriatide, amphotericin B, natural product, antifungal, potentiator, *Candida albicans*, ergosterol

## Abstract

Fungal cyclotetrapeptide nectriatide and its synthetic linear derivatives (nectriatide-based compounds, NCTs), having no antifungal activity, potentiated antifungal activity of amphotericin B (AmB) against *Candida albicans*. The mechanism of action was investigated using fluorescein-conjugated and biotin-tagged probes. Microscopy revealed fluorescent-probe localization at the *C. albicans* cell membrane. The biotinyl probe binding assay towards membrane lipids showed the highest affinity for ergosterol but no affinity for cholesterol. Ergosterol binding was confirmed by a liposome disruption assay. LC-MS quantification of AmB in *C. albicans* revealed that NCTs increased AmB binding to *C. albicans*. These results indicate that NCTs bind to ergosterol on the fungal plasma membrane, subsequently localizing AmB, explaining the observed potentiation of AmB fungicidal activity.

## Introduction

Fungal infections are estimated to impact an estimated 1.2 billion people worldwide ^1^. Unlike superficial infections, such as an athlete’s foot and dandruff, invasive fungal infections (IFIs) are deep-seated and include blood stream and systemic infections as well as infection of specific organs. Incidences of IFI appear to be increasing, with startlingly high estimated numbers of people affected; a recent report suggested an annual incidence of 6.5 million IFIs and 3.8 million deaths ^2^. Growing incidences of IFI are attributed to increasing immunocompromised populations; ageing, patients with HIV/AIDS, chronic lung diseases such as COPD, cancer and diabetes, and patients receiving immunosuppressive therapy. Additionally, new risks are emerging, including complications of patients with virus infection such as COVID-19 and influenza ^3^ ^4^. In clinical practice, four classes of antifungal drugs (azoles, echinocandins, polyenes, and pyrimidines) are used for IFI treatment. However, problems of side-effects and drug-drug interactions remain. And complicating matters, multi-drug-resistant *Candida auris* recently emerged as a cause of nosocomial outbreaks and complex infections^5^. Thus, new antifungal drugs are urgently needed, but a limited number of candidates are in the clinical development pipeline ^6^.

Amphotericin B (AmB) is a potent polyene macrolide, used to treat life-threatening IFIs for more than sixty years. AmB is an excellent antifungal due to its broad spectrum of activity, fungicidal efficacy, and low resistance rate. The antifungal activity is due to binding to ergosterol (Erg) in fungal cell membranes. However, as it also has an affinity to cholesterol (Chol) in mammalian cell membranes, AmB induces side-effects such as nephrotoxicity, nausea, fever, and vomiting, limiting its use. To overcome these limitations, new AmB formulations, including a lipid complex form ^7^ ^8^, liposomal form ^9^ ^10^, and nanoparticle form ^11^, are reported, with the AmB lipid complex (Abelcet) and liposomal AmB (AmBisome) forms finding clinical use. The Burke group recently reported reducing AmB derivative toxicity based on the analysis of AmB-Erg and AmB-Chol sponge complex ^12^.

To offset AmB side-effects, we propose a potentiator of antifungal AmB activity, to reduce AmB dosage. To achieve this, a screening of microorganism-derived AmB potentiators was performed, yielding several promising new compounds (**Supplemental Figure 1**). Simpotentin is a glycolipid comprising a sugar and C8 and C10 alkyl chains, achieved 4- to 8-fold potentiation in combination at 32 to 64 µg/mL ^13^. Phialotides with a long carbon chain (C20 ∼ C24) linked to one to three sugar moieties potentiated AmB activity by 16 to 32-fold at a lower dose (2.0 µg/mL) ^14^. Shodoamides are a tetradecadienamide structure with a triol butyl chain, and afforded a 4- to 8-fold potentiation in combination at 16 to 32 µg/mL ^15^. Unlike these amphipathic lipidic compounds, nectriatide (**1**), isolated from the culture broth of the fungus *Nectriaceae* sp. BF-0114, is a cyclic tetrapeptide composed of L-alanine, anthranilic acid, L-*N*-methyltyrosine, and L-valine ^16^ (**Figure 1**). Compound **1** potentiated AmB activity by 8-fold in combination at 32 µg/mL (**Table 1**). Recently, nectriatide derivatives were synthesized in the solution-phase and evaluated for AmB-potentiating activity ^17^. During this synthesis, linear intermediates (**2** and **3**, **Figure 1**) were found to exhibit greater AmB-potentiating activity than **1** with 8- to 32-fold potentiation in combination at 2.0 to 32 µg/mL. These findings prompted investigation into the mechanism of action. In this study, we demonstrate that nectriatide-based compounds (hereafter NCTs) including **2**, **3**, and chemical probes enhance AmB activity by binding cell membrane Erg.

**Fig. 1.**
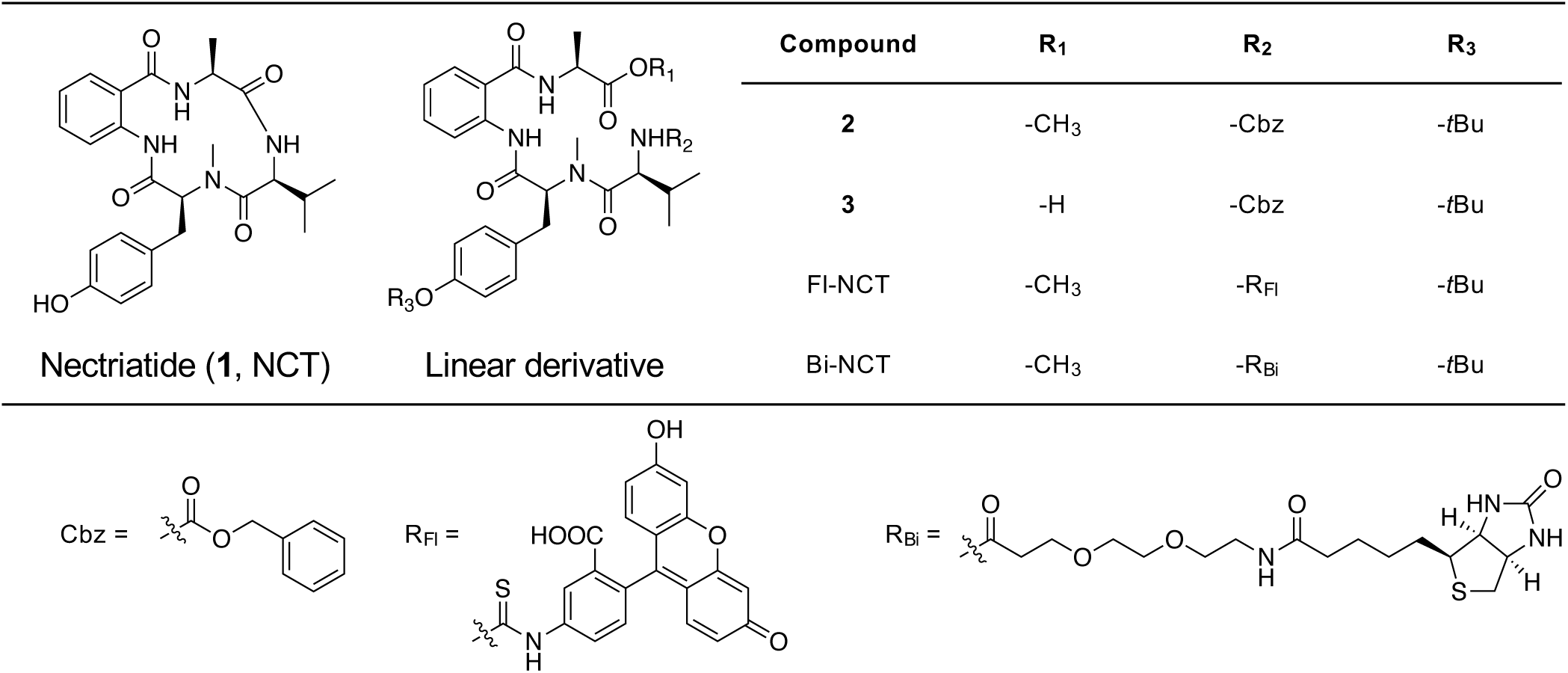
Structures of nectriatide (1) and the derivatives (NCTs).

**Table 1.**
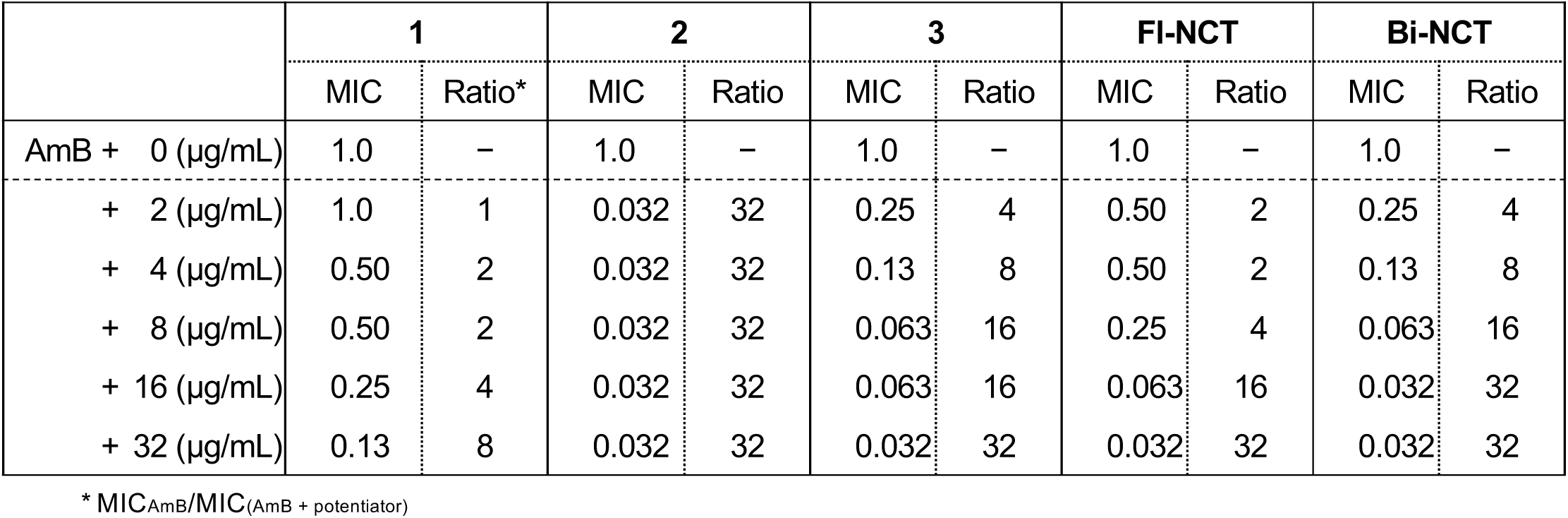
MIC values of AmB against *C. albicans* in the presence or absence of nectriatides.

|  | 1 |  | 2 |  | 3 |  | FI-NCT |  | Bi-NCT |  |
| --- | --- | --- | --- | --- | --- | --- | --- | --- | --- | --- |
|  | MIC | Ratio* | MIC | Ratio | MIC | Ratio | MIC | Ratio | MIC | Ratio |
| AmB + 0 (µg/mL) | 1.0 | – | 1.0 | – | 1.0 | – | 1.0 | – | 1.0 | – |
| + 2 (µg/mL) | 1.0 | 1 | 0.032 | 32 | 0.25 | 4 | 0.50 | 2 | 0.25 | 4 |
| + 4 (µg/mL) | 0.50 | 2 | 0.032 | 32 | 0.13 | 8 | 0.50 | 2 | 0.13 | 8 |
| + 8 (µg/mL) | 0.50 | 2 | 0.032 | 32 | 0.063 | 16 | 0.25 | 4 | 0.063 | 16 |
| + 16 (µg/mL) | 0.25 | 4 | 0.032 | 32 | 0.063 | 16 | 0.063 | 16 | 0.032 | 32 |
| + 32 (µg/mL) | 0.13 | 8 | 0.032 | 32 | 0.032 | 32 | 0.032 | 32 | 0.032 | 32 |

\* MIC<sub>AmB</sub>/MIC<sub>(AmB + potentiator)</sub>

## Results

### Synthesis of nectriatide chemical probes

Previously, nectriatide (**1**) and synthetic linear intermediates (**2** and **3**) were reported to potentiate AmB activity against *Candida albicans* ^17^. To investigate the mechanisms of action, fluorescent (Fl-NCT) and biotinylated derivatives (Bi-NCT) were synthesized (**Figure 1**, **Supplemental Scheme**). The AmB potentiating activity for each compound is noted in **Table 1**. Importantly, these linear intermediates more potently potentiated AmB activity than cyclic **1**. To determine whether these compounds enhance AmB cytotoxicity, a hemolysis assay was performed. AmB alone caused dose-dependently hemolysis with a HC_50_ value of 5.9 µg/mL (**Supplemental Figure 2**). However, NCTs alone exhibited no hemolysis even at 64 µg/mL. Interestingly, the co-administration of NCTs (32 µg/mL) with AmB did not enhance AmB hemolytic activity (HC_50_; 5.2 µg/mL in case of **2**) (**Supplemental Figure 2**). These results demonstrate specific AmB antifungal activity enhancement by NCTs.

### Localization of Fl-NCT in *C. albicans*

To visualize potentiator localization in *C. albicans*, Fl-NCT (64 µg/mL) was incubated with *C. albicans* at 35 °C for 1 h, then fluorescence intensity measured by fluorescence microscopy. Fl-NCT localization was observed in *C. albicans* plasma membranes and was more pronounced in the hyphal form than the yeast form (**Figure 2a**). Also, co-administration to *C. albicans* of a low concentration of AmB (a half of MIC, 0.50 µg/mL) further increased Fl-NCT plasma membrane localization (**Figure 2b**). In contrast, no fluorescence was observed by the inactive control potentiator, the fluorescently labeled L-valine analogue (**Supplemental Figure 3**).

**Fig. 2.**
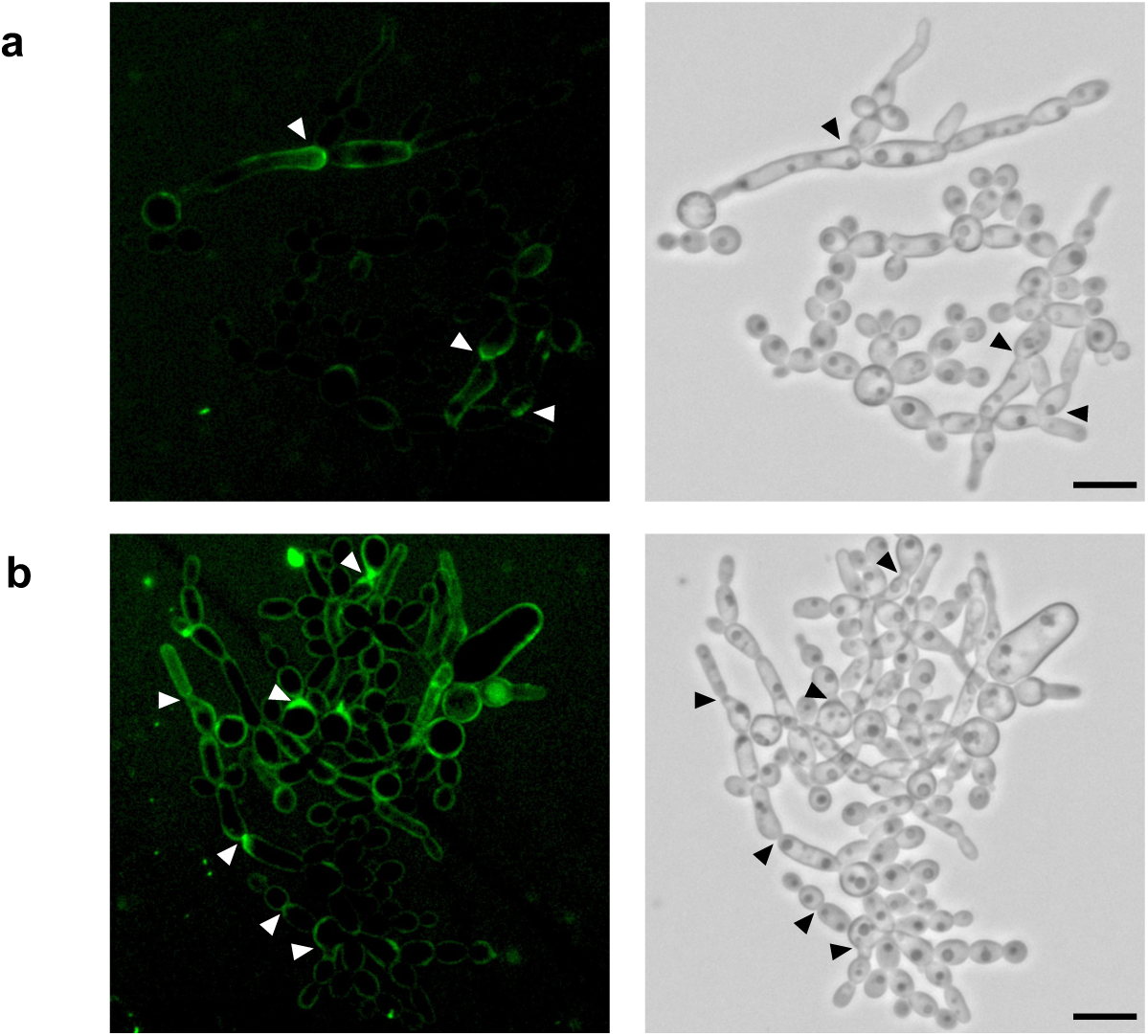
Localization of Fl-NCT on cell membrane in *C. albicans*. Cells were treated with (a) Fl-NCT (64 µg/mL) only or (b) Fl-NCT (64 µg/mL) combinated with AmB (0.50 µg/mL) at 35 °C for 1 h. Arrows represent higher fluorescence intensity area. Bars represent the scale of 10 µm.

### Binding of NCTs to fungal membrane lipids

NCT binding activity to the plasma membrane components [Erg, Chol, phosphatidylcholine (PC), phosphatidylethanolamine (PE), and phosphatidylinositol (PI)] was investigated using the Bi-NCT probe. Microplates coated with each of these lipids were incubated with Bi-NCT (64 µg/mL) for 1 h. After washing, the residual lipid-bound Bi-NCT was quantified by horseradish peroxidase (HRP)-conjugated streptavidin assay. As shown in **Figure 3**, Bi-NCT affinity for Erg was the strongest, followed by PI and PC. Importantly, no affinity towards Chol was observed for Bi-NCT, which suggests that NCTs selectively affect specific fungal membrane components. Since **2** and other NCTs did not enhance the hemolytic activity of AmB (**Supplemental Figure 2**), and PC and PI are common to mammals and fungi, Erg was expected to be the cellular target of NCTs.

**Fig. 3.**
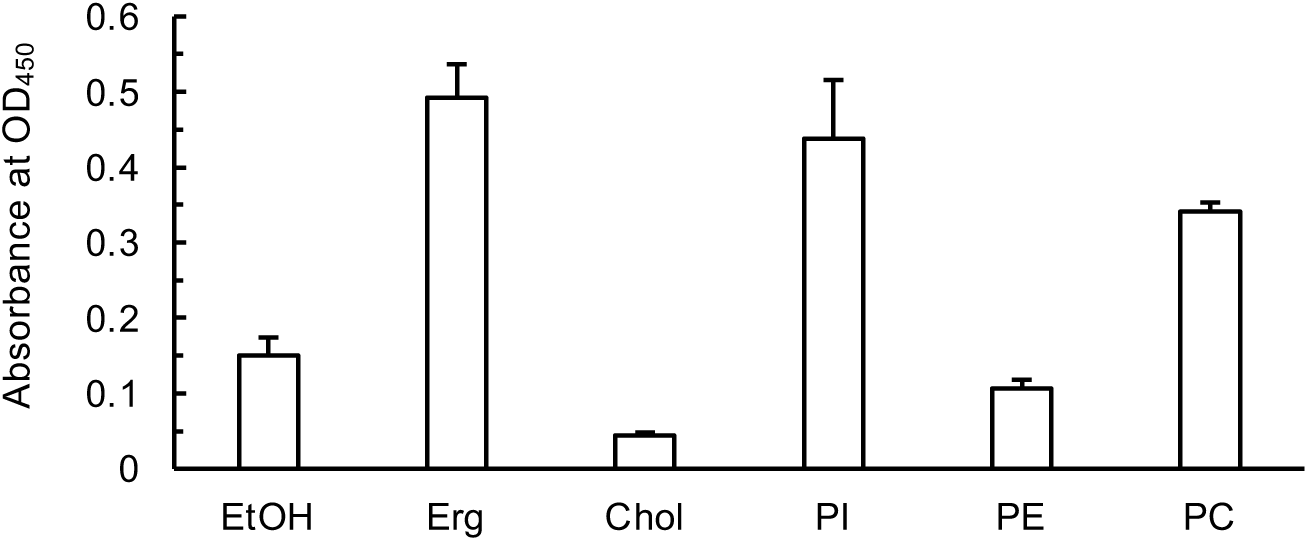
Affinity of Bi-NCT to ergosterol, cholesterol and membrane phospholipids. Data are presented as means ± SD from *n* = 3∼5 independent experiments.

Then, the affinity of **3** towards Erg-containing liposomes was evaluated. For a yeast cell membrane mimic liposome ^18^, composed of PC/PE/PI/Erg (5:4:1:2, w/w/w/w) and encapsulating fluorescent calcein, both AmB and **3** caused dose-dependent calcein leakage with EC_50_ values of 7.1 µg/mL and 52.2 µg/mL, respectively (**Figure 4a**). Furthermore, co-administration of **3** (8.0 µg/mL, a concentration causing no liposome disruption) with AmB induced a membrane damage at a lower concentration (EC_50_; 3.6 µg/mL) than AmB alone (7.1 µg/mL) (**Figure 4a**). Interestingly, Erg-free liposomes [PC/PE/PI (5:4:1, w/w/w)] were not disrupted by either AmB or **3** even at 64 µg/mL (**Figure 4b**). These findings demonstrate NCT binding to Erg, which increases the association of AmB to the liposome. Taken together, AmB and non-antifungal **3** disrupted Erg-containing liposomes, with a greater impact than individually.

**Fig. 4.**
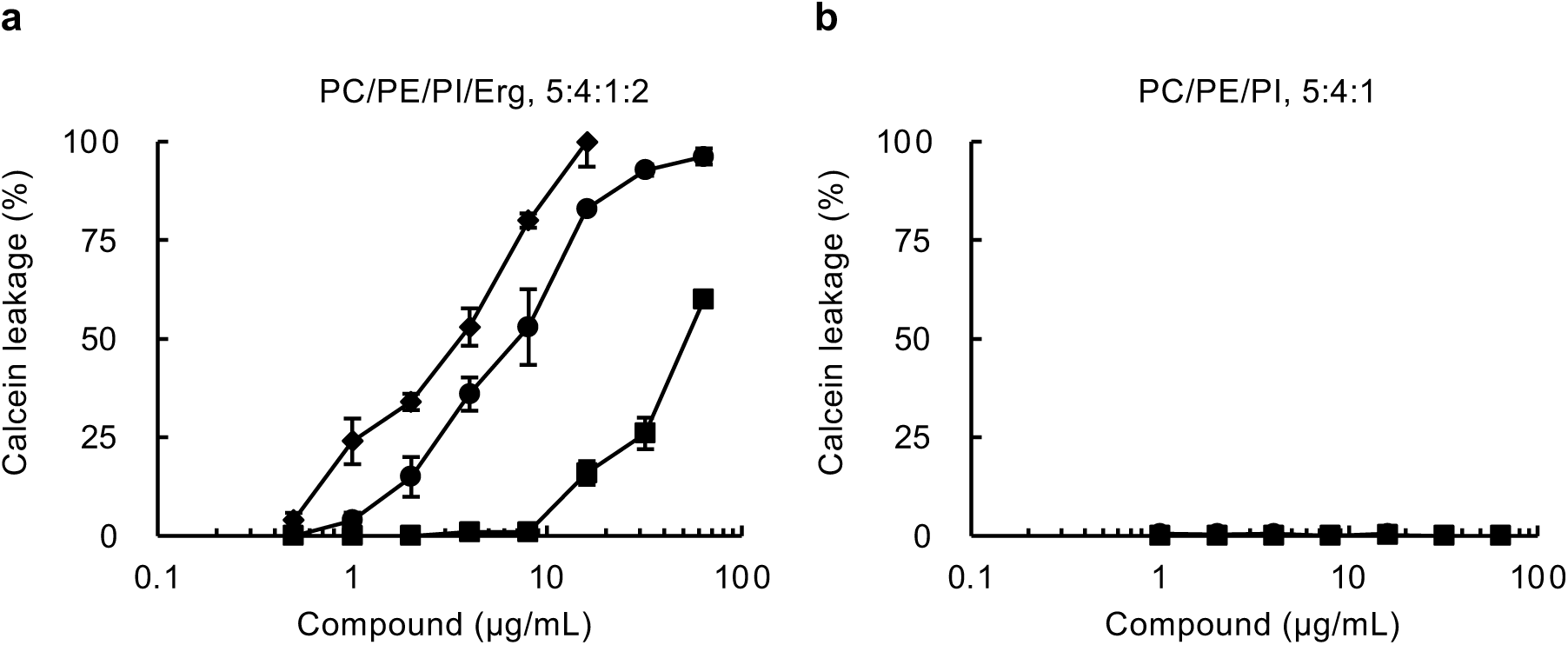
Affinity of 3 to liposomes containing ergosterol. (a) Calcein leakage from liposome [PC/PE/PI/Erg (5:4:1:2)] induced by AmB (circle), **3** (square) and combination with AmB and **3** (8 µg/mL) (diamond), (b) calcein leakage from liposome [PC/PE/PI (5:4:1)] induced by AmB (circle) and **3** (square). Data are presented as means ± SD from *n* = 3∼5 independent experiments.

### Recruit of AmB on *C. albicans* membrane by potentiators

The quantity of AmB bound to the fungal cell membrane in the presence or absence of potentiator **3** was measured by LC-MS analysis (**Supplemental Figure 4**). In this experiment, the incubation time of *C. albicans* with AmB was set to 1 h, as longer incubation times enable fungicidal activity by AmB. At AmB (0.063 mg/mL) alone, the amount of AmB bound to *C. albicans* was calculated to be 8.7 pg (**Figure 5**). The combination of **3** (0.050 to 12.5 µg/mL) increased AmB binding to the cells which plateaued (19.7 pg) at 3.1 µg/mL. These results show that NCTs bind Erg and recruit AmB to the cell membrane of *C. albicans*, potentiating AmB antifungal activity.

**Fig. 5.**
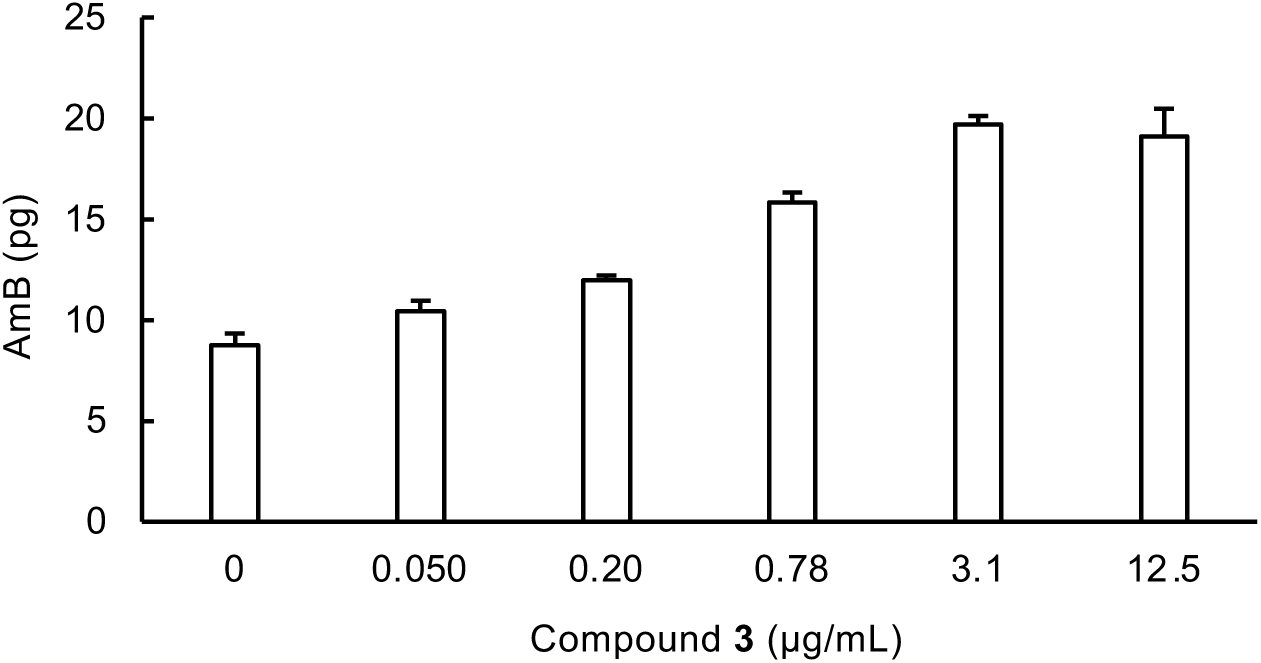
Quantification of AmB bound to *C. albicans* in the presence of 3 by LC-MS. Data are presented as means ± SD from *n* = 3∼5 independent experiments.

## Discussion

Nectriatide (**1**), a cyclotetrapeptide, was isolated from the culture broth of the fungus *Nectriaceae* sp. BF-0114 as a potentiator of AmB antifungal activity ^16^. During the synthetic study, its linear intermediates (**2** and **3**) were found to exhibit greater AmB-potentiating activity than **1** (**Figure 1**) ^17^. Importantly, these compounds exhibited no antifungal activity and no enhancement of AmB cytotoxicity to mammalian cells (**Table 1**, **Supplemental Figure 2**). To study the mechanism of action, Fl-NCT and Bi-NCT were synthesized. From fluorescent microscopy analysis, Fl-NCT was observed to localize in *C. albicans* membranes, and more densely in the hyphal form (**Figure 2a**). *C. albicans* shows yeast and hyphae dimorphism, adapting to a variety of microenvironments ^19^. Upon hyphal extension, Erg is enriched at the bud tip during bud emergence and at the apex during germ tube emergence ^20^, which explains stronger fluorescence responses of Fl-NCT for the hyphal form. Accordingly, interactions between Bi-NCT and cell membrane lipids were investigated by binding assays. Fl-NCT was first used, but its high lipophilicity resulted in localization within the assay plate. Therefore, the less lipophilic Bi-NCT was used. From these assays, Bi-NCT presented the highest affinity towards Erg, but yielded no affinity towards Chol (**Figure 3**), which may explain the lack of AmB cytotoxicity enhancement by NCTs towards mammalian cells. For the more sensitive liposome assay, **3** was selected as it has a lower lipophilicity than **2**. Both AmB and **3** disrupted liposomes composed of PC/PE/PI/Erg (5:4:1:2, w/w/w/w), but did not damage Erg-free liposomes [PC/PE/PI (5:4:1, w/w/w)] (**Figure 4**). These results reflect those of the direct lipid binding assays. Intriguingly, co-administration of AmB with **3** (8.0 µg/mL), a concentration at which liposomes are normally not disrupted, induced liposome disruption (**Figure 4a**). Additionally, potentiator **3** was found to recruit AmB in *C. albicans* in a dose-dependent manner (0.05 to 3.1 µg/mL) (**Figure 5**). These findings indicate that NCTs bind to Erg and increase AmB accumulation on *C. albicans* membranes, leading to AmB fungicidal activity potentiation. The reason for the plateau observed for AmB binding is proposed due to co-treatment of AmB with 3.1 µg/mL of **3** was sufficient to completely disrupt the cell membrane, thus preventing further AmB binding.

Two models are proposed for the mechanism of action of AmB. In the barrel-stave model, AmB forms transmembrane pores with Erg in fungal cell membranes, inducing the collapse of vital ion gradients, thereby killing the fungal cells ^21^ ^22^. Umegawa et al. recently reported AmB assembly into heptameric molecular ion channels observed by solid-state NMR analysis ^23^. In the sterol sponge model, extra-membranous aggregates of AmB physically extract Erg from the fungal membrane, thereby killing the fungal cells without pore formation ^24^ ^25^. We speculate that NCTs may form more/bigger pores in *C. albicans* membranes or extract more Erg from *C. albicans* membranes by accumulating more AmB on *C. albicans* membranes.

Many natural products are known to target cell membrane lipids ^26^. Detergents such as biosurfactants and saponins interact with a variety of lipids. On the other hand, microbial polyene macrolides, including AmB, target sterols. Theonellamide targets 3β-hydroxysterols ^27^, papuamide targets phosphatidylserine ^28^, and both cinnamycin and duramycin target PE ^29^. However, these compounds induce cell membrane damage, leading to cytotoxicity, antibacterial activity, or/and antifungal activities. Importantly, NCTs exhibited no antifungal activity despite their affinity to Erg.

In summary, non-antifungal NCTs bind to Erg, increasing AmB recruitment to the plasma membrane of *C. albicans*, leading to AmB fungicidal activity potentiation. How NCTs recognize different sterols within the cell membrane remains elusive. Study of the mechanism of action and *in vivo* efficacy of NCTs is ongoing.

## Material and methods

### General

NMR spectra were obtained using an NMR System 400 MHz spectrometer (DD2-400NB; Agilent Technologies, Santa Clara, CA, USA). MS analyses were performed using the Xevo G2-XS QTof (Waters, Milford, MA, USA). HPLC was performed using Prominence systems (Shimadzu, Kyoto, Japan) equipped with the DGU-20A_5_ degasser, the SPD-20A UV detector, and the LC-20AT pump.

Absorbance intensity was measured using a microplate absorbance reader (iMark; Bio-Rad, Hercules, CA, USA). Fluorescence intensity was measured using a multi-mode microplate reader (SpectraMax M5; Molecular Devices, Sano Jose, CA, USA). Microscopic observation was performed using a fluorescence microscope BZ-710 (Keyence, Osaka, Japan).

### Materials

Organic solvents were as follows. DMSO, MeOH, EtOH, MeCN and CHCl_3_ were purchased from Nacalai-tesque (Kyoto, Japan). *N*,*N*-Dimethylformamide (DMF) and *N*,*N*-Diisopropylethylamine (DIPEA) were purchased from Watanabe Chemical Industries (Hiroshima, Japan).

Materials used for AmB potentiating activity assays were as follows. Amphotericin B was purchased from Sigma-Aldrich (now Merck, Darmstadt, Germany). RPMI 1640 medium dry powder was purchased from Thermo Fisher Scientific (Waltham, MA, USA). 3-Morpholinopropanesulphonic acid (MOPS) was purchased from DOJINDO (Kumamoto, Japan).

Materials used for lipid binding and liposome assays were as follows. A microtiter plate (Immulon 1B) was purchased from Thermo Fisher Scientific. 2-Amino-2-hydroxyl-1,3-propanediol (Tris), ergosterol (Erg), Tween 20, NaCl, hydrochloric acid, and skim milk were purchased from FUJIFILM Wako Pure Chemical. Cholesterol (Chol), L-α-phosphatidylcholine (PC) (from egg yolk), L-α-phosphatidylethanolamine (PE) (from egg yolk), phosphatidylinositol (PI) (from soy), and Triton X-100 were purchased from Merck. 3,3’5,5’-Tetramethylbenzidine (TMB) solution for ELISA, horseradish peroxidase (HRP) conjugated streptavidin, *N*-hydroxysuccinimide (NHS)-PEG_2_-biotin, and L-valine methyl ester hydrochloride were purchased from Tokyo Chemical Industry (Tokyo, Japan). Calcein and fluorescein isothiocyanate isomer-I (FITC-I) were purchased from DOJINDO.

### Assay for potentiation of antifungal AmB activity

The broth microdilution method using a 96-well microplate (AS ONE, Osaka, Japan) was performed according to the guidelines of the Clinical Laboratory Standards Institute (CLSI) documents M27-A4 ^30^ and our established method ^14, 16^. Five colonies of *C. albicans* ATCC 90029 strain with diameters of 1 mm were suspended in sterile saline (0.85% NaCl) to adjust to a 0.5 McFarland standard by spectrophotometric measurement. The seed of yeast was diluted 2,000 times with RPMI 1640 medium (1.04% RPMI 1640 medium dry powder and 3.45% MOPS, pH 7.0 adjusted by 2 M NaOH). The diluted seed (180 µL) and RPMI 1640 medium (18 µL) were added to each well of a 96-well microplate with or without serial concentrations of test compounds (each of 1 µL, DMSO solution). The 96-well plate was incubated at 35 °C for 24 h. After incubation, the OD_550_ value of each well was measured by a microplate reader (iMark) to assess the MIC. The test compound antifungal activity was evaluated at concentrations ranging between 0 and 64 µg/mL. Antifungal AmB potentiation activity was tested in a combination with the test compound (2.0 to 32 µg/mL). MIC was defined as the lowest concentration of AmB at which fungal growth was inhibited by 90% from control growth (no AmB).

### Assay for hemolytic activity

The assay for hemolytic activity was performed as previously described ^31^, with a slight modification. Briefly, sheep preserved blood (Japan Bio Serum, Hiroshima, Japan) was inoculated into a 50 mL centrifuge tube, and the tube centrifuged at 500 *g* for 5 min at 4 °C. The supernatant was discarded, then cells suspended in sterile saline and washed four times by a centrifugation under the same described conditions. After washing, cells were adjusted to 7.0 × 10^7^ cells/mL using sterile saline. The suspension (99 µL) was transferred to each well of a round-bottom 96-well microplate, then a test sample (1 µL) was added to each well. After incubation for 2 h at 37 °C, the plate was centrifuged at 3,000 rpm for 15 min at 4 °C. The supernatant (80 µL) was transferred to a 96-well plate, then the color intensity of each well was measured with a microplate reader (iMark), reading the absorption at 550 nm. To determine 100% hemolysis, 0.1% Triton X-100 was added to the red blood cells. The percentage of hemolysis caused by a compound was calculated as follows: hemolysis (%) = 100 × (H - H_0_) / (H_t_ - H_0_), where H represents the hemolysis achieved after addition of a compound, and H_0_ and H_t_ represent the hemolysis with only DMSO and TritonX-100, respectively. The HC_50_ value was defined as the lowest concentration of a test compound at which hemolysis reached 50%.

### Fluorescent microscopy analysis

A *C. albicans* colony was inoculated into a 50 mL centrifuge tube containing 10 mL RPMI 1640 medium, and the tube shaken on a rotary shaker (180 rpm) for 16 h at 35 °C. After centrifugation, the cells were suspended in RPMI 1640 medium to adjust to the OD_600_ value of 0.32 using a microplate reader. The suspension (300 µL) was transferred to each well of a 48-well microplate (Sumitomo Bakelite, Tokyo, Japan), then a test sample (3 µL) was added to each well. After incubation for 1 h at 35 °C, each culture was transferred to a 1.5 mL microtube, then the cells washed three times with 200 µL of phosphate-buffered saline (PBS) by flash centrifugation. The cell suspension prepared with 5 µL of PBS was spread on a glass slide (Matsunami Glass Industry, Osaka, Japan). After the slide was covered with a cover glass (Matsunami Glass Industry), fluorescence microscopy was performed using a fluorescent microscope (BZ-710) equipped with Plan Apochromat 40× and 100× lenses (Nikon Instruments, Melville, NY, USA).

### Lipid binding assay

The lipid binding assay was performed as described previously ^27^ ^32^, with a slight modification. Briefly, plate wells (Immulon 1B) were coated with 20 µL of lipid solution (50 µM) in EtOH by evaporation over 2 h at room temperature. After blocking the wells with 100 µL of Tris-buffered saline (10 mM Tris·HCl, pH 7.4, 150 mM NaCl) containing 5.0% (w/v) skim milk (blocking buffer) at 4 °C for 16 h, wells were incubated with Bi-NCT (64 µg/mL) in blocking buffer for 1 h at room temperature. After washing the wells three times with 100 µL of PBS containing 0.02% tween 20, the wells were incubated with 100 µL of HRP-conjugated streptavidin diluted 2,000 times in PBS for 1 h at 35 ℃. After washing the wells three times with PBS, the wells were incubated with 100 µL of TMB solution at room temperature for 30 min. Finally, the wells were incubated with 100 µL of 1 M HCl at room temperature for 1 h. The color intensity of each well was measured with a microplate reader (iMark), reading the absorption at 450 nm.

### Liposome assay

Liposomes mimicking the yeast cell membrane ^18^ were prepared as follows. Aliquots of lipid stock solutions containing 1.0 mg PC, 2.4 mg PE, 0.60 mg PI and 1.2 mg Erg dissolved in CHCl_3_ (PC/PE/PI/Erg, 5:4:1:2, w/w/w/w) were placed in a flask, and evaporated under vacuum to provide a thin homogenous lipid film for 16 h at 25 °C. The lipid was then dissolved in 1.0 mL of Tris-buffered saline containing 0.20 mg/mL calcein. The mixture was heated at 70 °C for a few seconds, and then stirred by mixer for 30 seconds, with this repeated five times. The resultant suspension was extruded 21 times through 100 µm-polycarbonate filters (Cytiva, Tokyo, Japan) with an extruder (Avanti polar lipid, Alabaster, AI, USA). Unencapsulated calcein was removed with filtration using 100 kDa-cut off Amicon Ultra (Merck) by centrifugation for 10 min at 14,000 *g*, and concentrated liposomes (approximately 25 µL) were collected. The liposomes were diluted 300-fold with Tris-buffered saline, 99 µL of the suspension was added to each well of a 96-well black microplate (Corning, Corning, NY, USA) with 1 µL of a test compound in DMSO or DMSO. After incubation for 50 min at room temperature, liposome calcein leakage was monitored by measuring fluorescence intensity at 515 nm using an excitation wavelength of 495 nm with a multi-mode microplate reader (SpectraMax M5). To determine 100% calcein release, 0.1% Triton X-100 was added to the liposomes. The percentage of calcein leakage caused by the compounds was calculated as follows: calcein leakage (%) = 100 × (F - F_0_) / (F_t_ - F_0_), where F represents the fluorescence intensity achieved after compound addition with F_0_ and F_t_ representing fluorescence intensities obtained with only DMSO and Triton X-100, respectively. The liposomes (PC/PE/PI, 5:4:1, w/w/w) were also prepared in the same procedure described above. The EC_50_ value was defined as the lowest concentration of a test compound at which 50% calcein leakage occurred.

### Quantification of AmB bound to *C. albicans* by LC-MS analysis

A colony of *C. albicans* was inoculated into a 50 mL centrifuge tube containing 10 mL RPMI 1640 medium, and the tube shaken on a rotary shaker (180 rpm) for 16 h at 35 °C. After centrifugation, the cells were suspended in RPMI 1640 medium and OD_600_ value adjusted to 0.27 indicated by microplate reader (iMark). The suspension (200 µL) was transferred to each well of a 96-well microplate, the cells were treated with AmB (0.063 µg/mL) and **3** (0 to 12.5 µg/mL) for 1 h at 35 °C. After each suspension was transferred to a 1.5 mL microtube, the supernatant containing free AmB was removed by a centrifugation at 12,000 *g* for 5 min. The precipitated cells were treated with 200 µL of MeOH and sonicated for 1 h to extract AmB bound to *C. albicans*. The MeOH solution was subjected to LC-MS analysis [column, Acqulty i.d. 2.1 × 50 mm (Waters); mobile phase, 5-min linear gradient from 5.0 to 100% MeCN with 5.0 mM ammonium acetate aqueous solution; flow rate, 0.30 mL/min; column oven, 40 °C] with a MS spectrometer. Under these conditions, AmB was detected with a retention time of 3.7 min.

### Synthesis of nectriatide chemical probes

To remove the Cbz group of **2**, a flask containing **2** (50 mg, 72.6 µmol) and 10% Pd/C (25 mg) in EtOH (1 mL) was evacuated and filled with hydrogen gas. The mixture was then stirred vigorously for 4 h at room temperature. After that, the mixture was filtered through a Celite pad and the filtrate concentrated. The primary amine residue was used for next reactions without further purification because of its instability.

The synthesis of Fl-NCT was carried out as follows. A glass vial containing the primary amine residue (1.3 mg, 2.3 µmol) and FITC-I (1.8 mg, 4.6 µmol) in DMF (0.4 mL) was stirred for 2 h at room temperature under protection from light. After that, the solvent was evaporated. The residue was purified by preparative HPLC [Column; PEGASIL ODS C18 SP100 i.d. 20 × 250 mm (Senshu Scientific, now Kitahama, Osaka, Japan), mobile phase, 68% MeCN aq isocratic; flow rate, 8.0 mL/min; detection, UV at 210 nm]. Under these conditions, Fl-NCT was eluted with a retention time of 11 min. The peak was collected and concentrated to dryness yielding Fl-NCT (1.2 mg, 1.3 µmol, 57%). ESI-MS: calcd for C_51_H_54_N_5_O_11_S [M+H]^+^ 944.3542, found 944.3624.

The synthesis of Bi-NCT was carried out as follows. A glass vial containing the primary amine residue (2.0 mg, 3.4 µmol) and NHS-PEG_2_-biotin (2.0 mg, 4.0 µmol) in CH_2_Cl_2_ (0.4 mL) was stirred for 2.5 h at room temperature. After that, the solvent was evaporated. The residue was purified by preparative HPLC [Column; PEGASIL ODS C18 SP100 i.d. 20 × 250 mm, mobile phase, 50% MeCN aq isocratic; flow rate, 8.0 mL/min; detection, UV at 210 nm]. Under these conditions, Bi-NCT was eluted with a retention time of 12 min. The peak was collected and concentrated to dryness yielding Bi-NCT (1.4 mg, 1.5 µmol, 44%). ESI-MS: calcd for C_47_H_70_N_7_O_11_S [M+H]^+^ 940.4854, found 940.5830.

The synthesis of FITC conjugated valine analogue was performed as follows. A glass vial containing L-valine methyl ester hydrochloride (1.25 mg, 9.5 µmol) and FITC-I (5.0 mg, 12.8 µmol) in DMF (1.5 mL) with DIPEA (7.0 µL) was stirred for 2 h at room temperature under protection from light. After that, the reaction mixture was purified by preparative HPLC [Column; PEGASIL ODS C18 SP100 i.d. 20 × 250 mm mobile phase, 50% MeCN aq containing 0.05% TFA isocratic; flow rate, 6.0 mL/min; detection, UV at 210 nm]. Under these conditions, FITC conjugated valine analogue was eluted with a retention time of 18 min. The peak was collected and concentrated to dryness yielding FITC conjugated valine analogue (2.9 mg, 5.9 µmol, 63%). ESI-MS: calcd for C_26_H_21_N_2_O_6_S [M+H]^+^ 489.1120, found 489.1113.

## Acknowledgements

We thank Ms. Noriko Sato (Graduate School of Pharmaceutical Sciences, Kitasato University) for NMR measurements. This work was financially supported by JSPS KAKENHI Grant numbers JP19K05719 (KN), JP21K15284 (KK), and JP22K05333 (KN), The Tokyo Biochemical Research Foundation (now Chugai Foundation for Innovative Drug Discovery Science: C-FINDs) (KK), The Research Foundation for Pharmaceutical Sciences (KK) and Kitasato University Research Grant for Young Researchers (KK).

## Compliance with ethical standards

## Supplementary information

**Supplemental Scheme.**
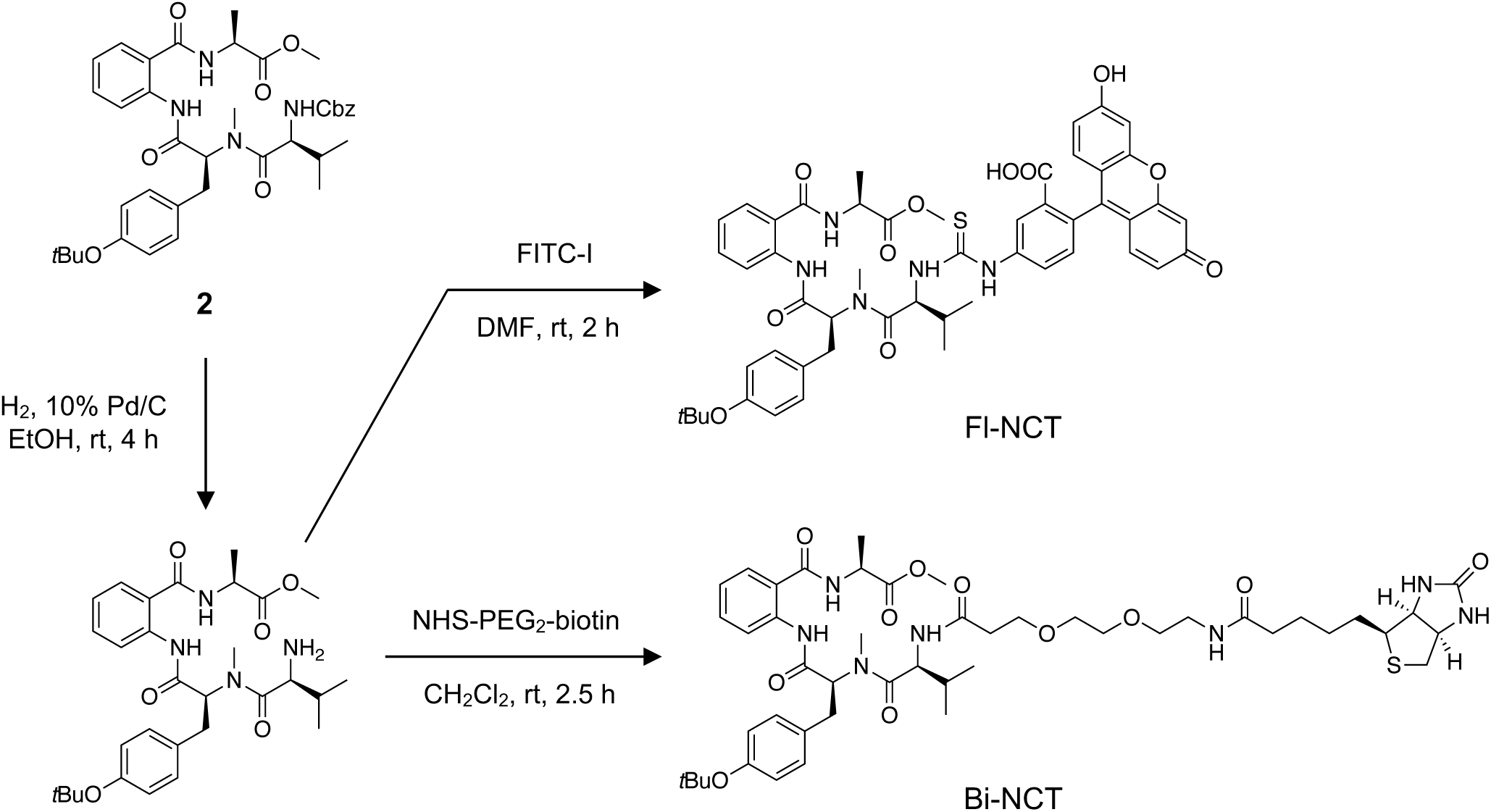
The synthesis of nectriatide chemical probes.

**Supplemental Figure 1.**
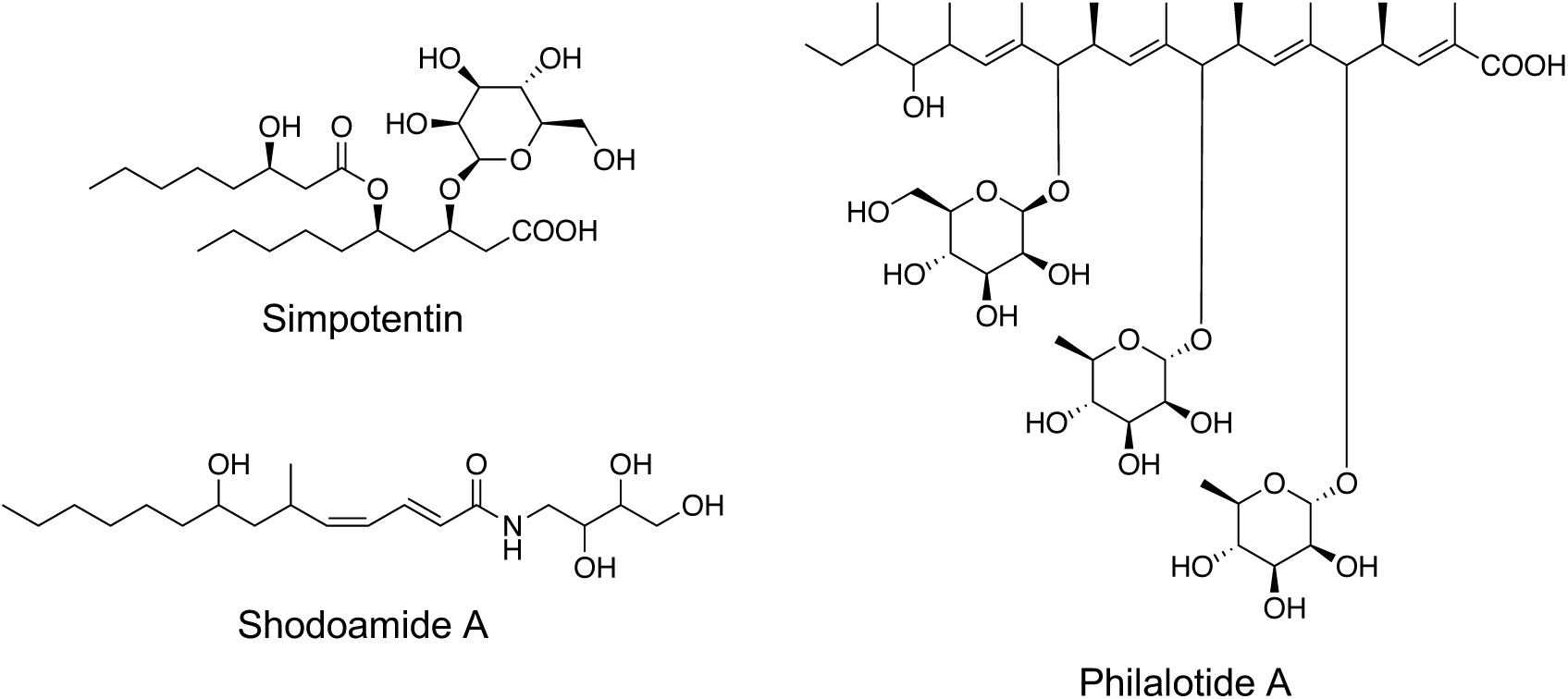
AmB potentiators discovered from microorganisms.

**Supplemental Figure 2.**
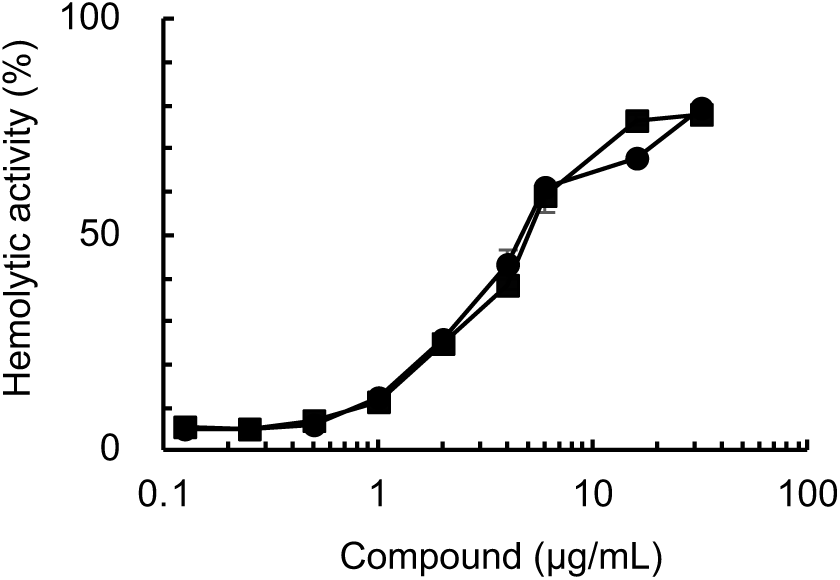
Hemolytic activity of AmB in the presence of **2**. Hemolysis induced by AmB (circle) and combination with AmB and **2** (32 µg/mL) (square). Data are presented as means ± SD from *n* = 3 independent experiments.

**Supplemental Figure 3.**
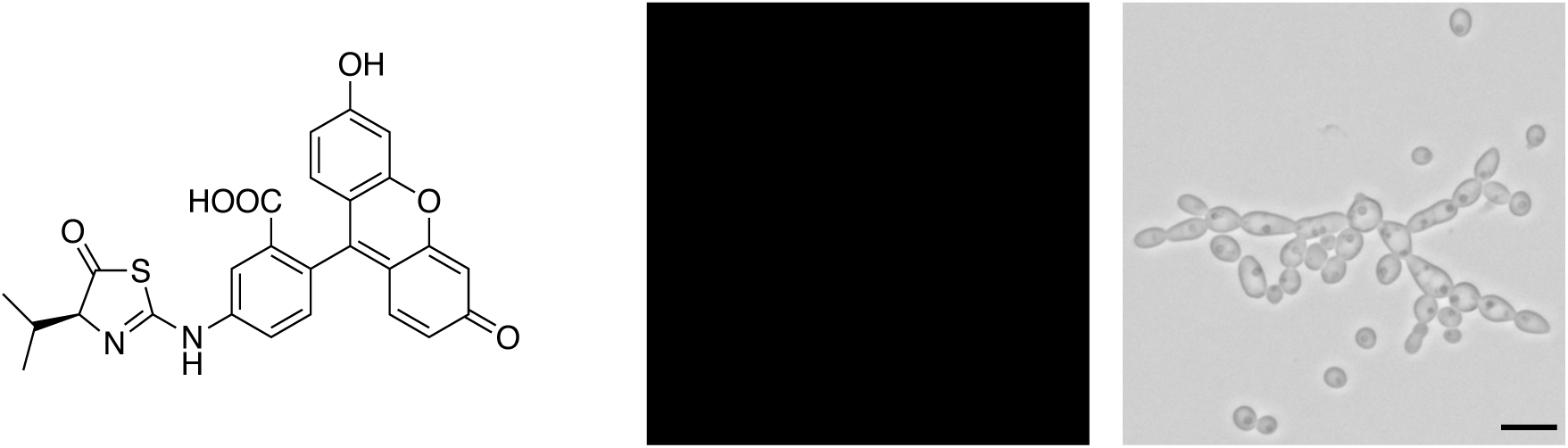
No localization of FITC conjugated L-valine analogue on cell membrane in *C. albicans*. Cells were treated with FITC conjugated L-valine analogue (64 µg/mL) at 35 °C for 1 h, then fluorescence intensity measured by fluorescence microscopy. Bar represents the scale of 10 µm.

**Supplemental Figure 4.**
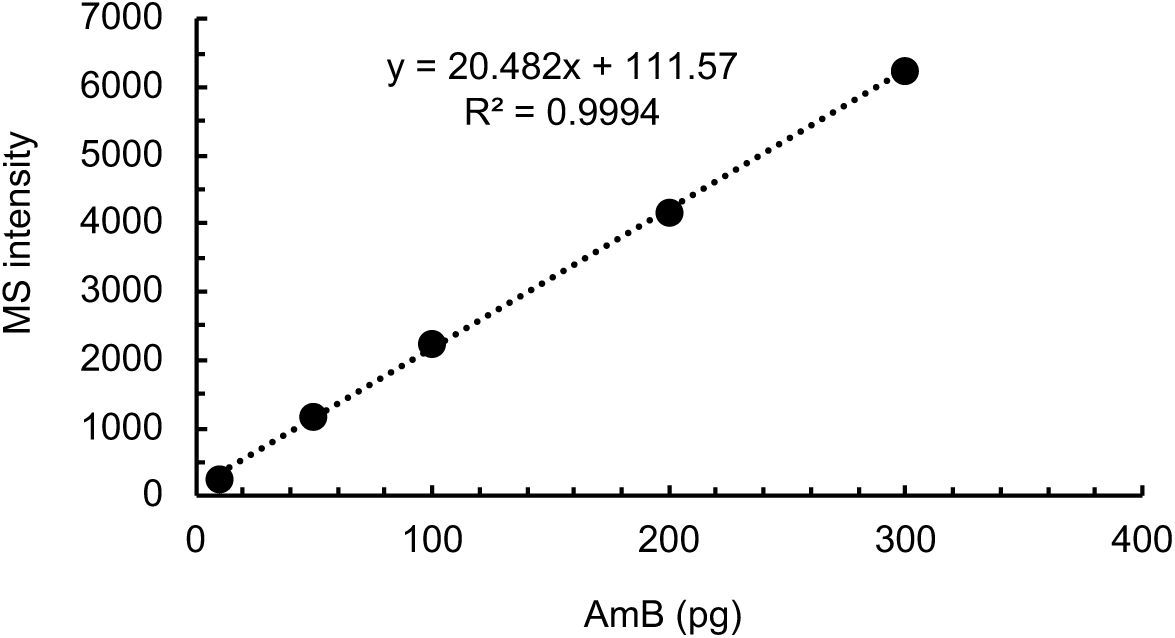
Calibration curve between the amount of AmB and MS intensity

**Supplemental Figure 5.**
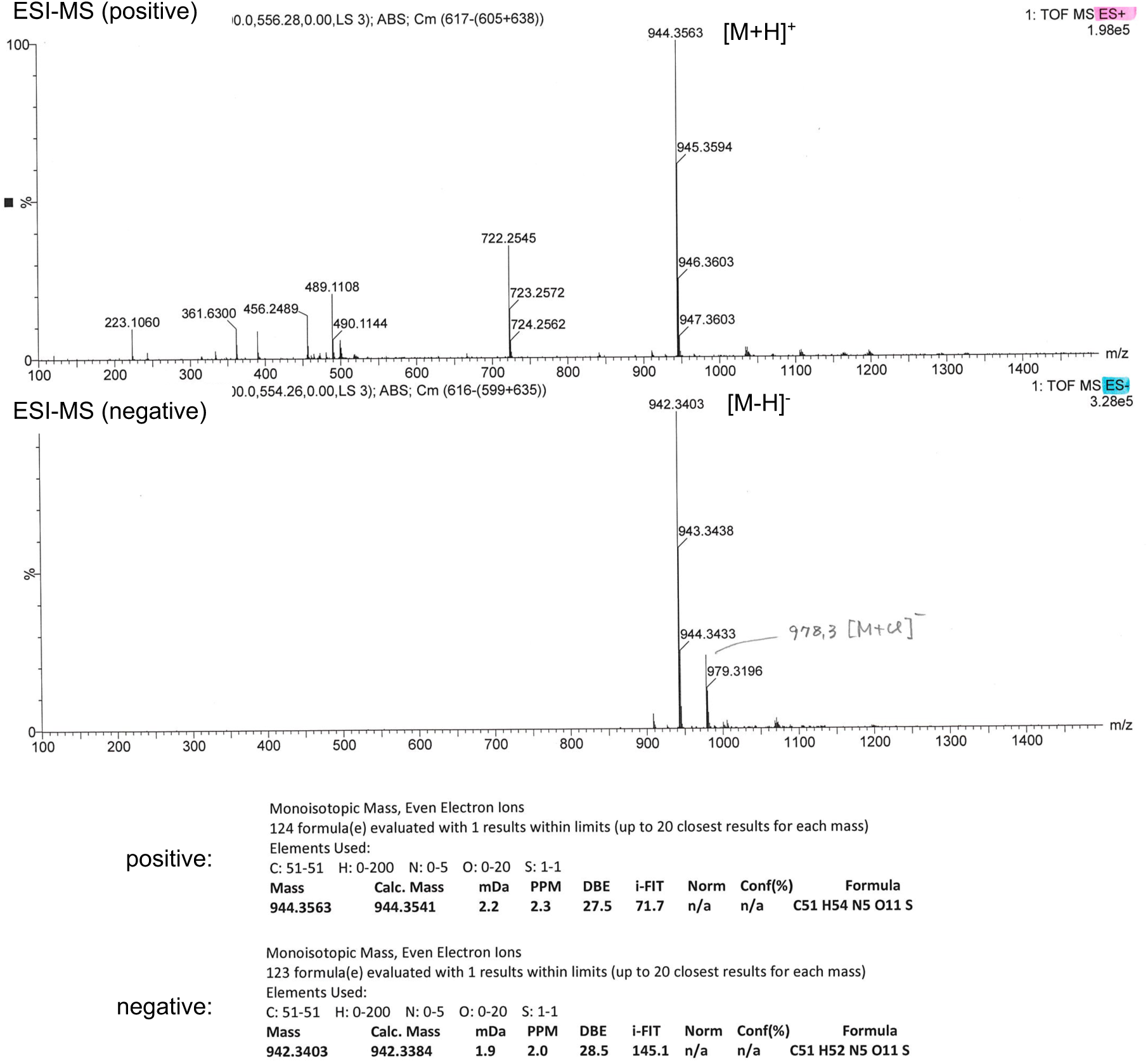
ESI-MS spectra of Fl-NCT.

**Supplemental Figure 6.**
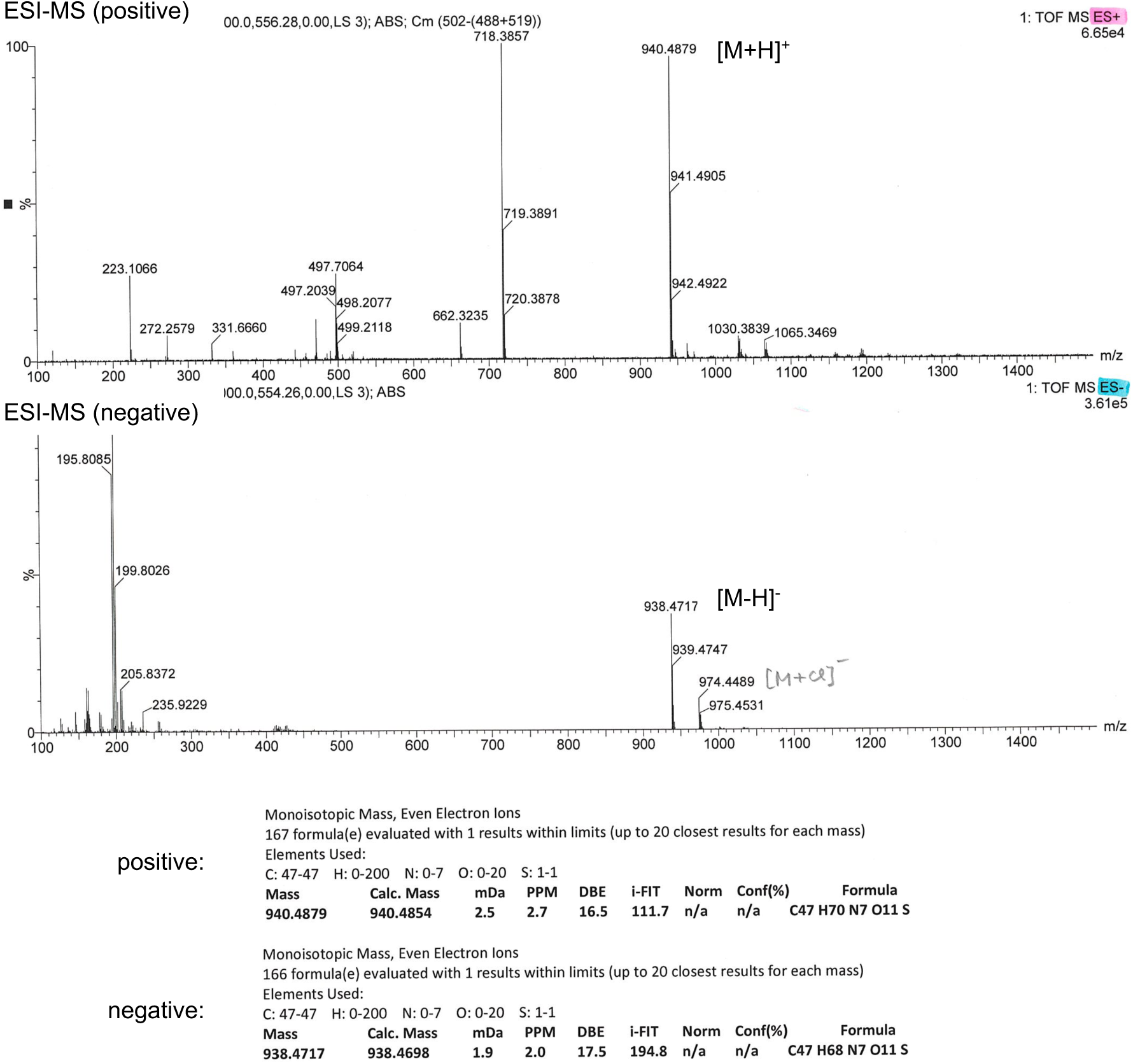
ESI-MS spectra of Bi-NCT.

**Supplemental Figure 7.**
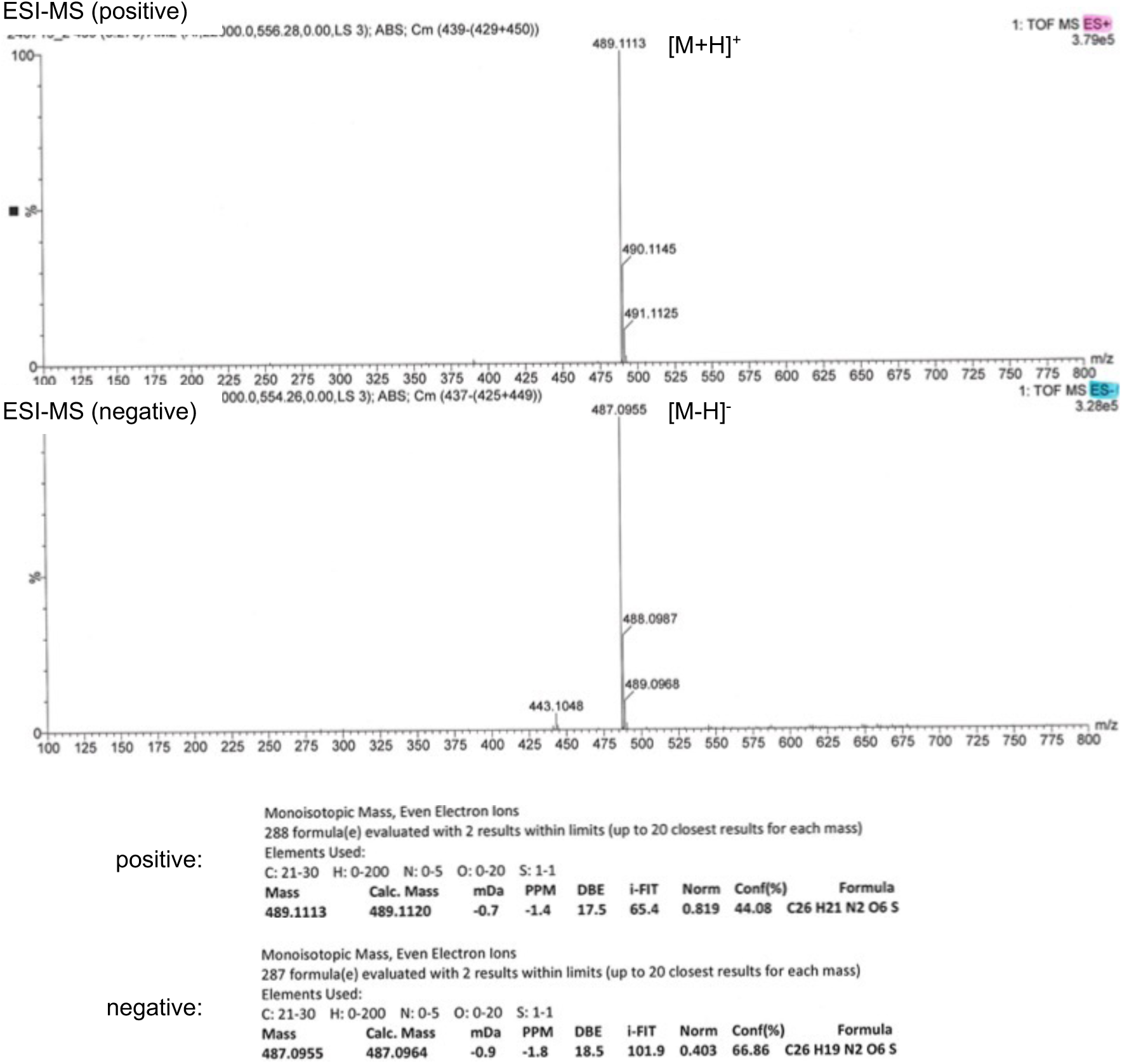
ESI-MS spectra of fluorescently labeled L-valine methyl ester analogue.

